# Polygenic predictors of age-related decline in cognitive ability

**DOI:** 10.1101/375691

**Authors:** Stuart J. Ritchie, W. David Hill, Riccardo E. Marioni, Gail Davies, Saskia P. Hagenaars, Sarah E. Harris, Simon R. Cox, Adele M. Taylor, Janie Corley, Alison Pattie, Paul Redmond, John M. Starr, Ian J. Deary

## Abstract

Polygenic scores can be used to distil the knowledge gained in genome-wide association studies for prediction of health, lifestyle, and environmental factors in independent samples. In this preregistered study, we used fourteen polygenic scores to predict variation in cognitive ability level at age 70 and cognitive change from age 70 to age 79 in the longitudinal Lothian Birth Cohort 1936 study. The polygenic scores were created for phenotypes that have been suggested as risk or protective factors for cognitive ageing. Cognitive abilities within old age were indexed using a latent general factor estimated from thirteen varied cognitive tests taken at four waves, each three years apart (initial *n* = 1,091 age 70; final *n* = 550 age 79). The general factor indexed over two-thirds of the variance in longitudinal cognitive change. We also ran an additional analysis using an age-11 intelligence test to index cognitive change from age 11 to age 70. Several polygenic scores were associated with the level of cognitive ability at age-70 baseline (range of standardized *β*-values = –178 to .264), and the score for education was associated with cognitive change from childhood to age 70 (standardized β = .102). None was statistically significantly associated with variation in cognitive change between ages 70 and 79. *APOE* e4 status made a significant prediction of cognitive decline from age 70 to 79 (standardized *β* = –319 for carriers vs. non-carriers). The results suggest that the predictive validity for cognitive ageing of polygenic scores derived from genome-wide association study summary statistics is not yet on a par with *APOE* e4, a more well-established predictor.

## Introduction

Mean levels of cognitive function decline as people grow older, even in those without dementia. This affects many important cognitive functions, such as memory, processing speed, and reasoning ability^1,2,3^, with so-called “crystallized” abilities, such as vocabulary, less affected. There is strong evidence that declines across all abilities are correlated: cognitive ageing, as with individual differences in cognitive ability level, is substantially a general phenomenon^4,5,6^. Declines in cognitive abilities in older age have practical consequences for daily life and independent living: they have been linked to lower ability to perform everyday functions such as understanding medicine labels^7^, and to increased vulnerability to financial fraud^8,9^. Discovering predictors of variation in cognitive ageing might help us to identify those at highest risk of more rapid decline, and—to the extent that such predictors are confirmed to be causal—devise appropriate interventions. In the present study, we assessed the value of a panel of genetic risk scores in predicting variation in general cognitive decline in a generally healthy sample across the eighth decade of life.

Many studies have investigated whether variables that are known to correlate cross-sectionally with cognitive ability are also predictive of variation in its decline. Numerous such factors have been tested, but few have been replicated consistently^10^. For instance, although higher educational attainment has been found to be predictive of shallower rates of cognitive decline^11,12^—a finding which has informed theories of “cognitive reserve”^13^—other studies have not found this same effect^14,15^. Other potential predictors, with varying degrees of evidentiary support, include physical fitness, as measured by variables such as grip strength and lung function^16,17^ (see ref.^18^ for a review), personality traits such as conscientiousness^19^ (see ref.^20^ for a review), and type 2 diabetes^21^ (see ref. for a review).

Here, we investigate potential genetic predictors of cognitive level at age 70, and relative cognitive decline from age 11 to 70 years and from age 70 to 79 years. One such predictor is well-known already: carriers of either one or two *APOE* e4 alleles (as opposed to no such alleles) are not just at higher risk of a diagnosis of Alzheimer’s disease^23,24^, but also appear to be at risk of steeper cognitive decline^25^. In recent years, however, a new method has become commonly used in research investigating genetic prediction of traits: polygenic scoring. This method uses summary data from published genome-wide association studies (GWAS) that have tested the correlations of millions of single-nucleotide polymorphisms (SNPs) with phenotypes of interest. Using the weightings (regression coefficients) for each SNP from these data, genotyped individuals in an independent sample (one not included in the original GWAS) can have a polygenic score (PGS) calculated that indexes their genetic liability to a certain disease, or their probability of a higher level of a particular trait^26^. Through metaanalysis, and through the collection of ever-larger datasets, the sample size, and thus the statistical power, of GWAS studies continues to increase. For example, the variance explained in educational attainment in independent samples by the educational attainment PGS has increased alongside the sample size of the discovery GWASs^27,28^.

PGSs can be used to predict variables other than their “own” phenotype. The PGS for educational attainment, for example, has been shown not just to predict educational attainment but also, among others, cognitive ability^29^, social mobility^30,31^, and longevity^32^. It is possible, then, that the genetic variants linked to phenotypic predictors of cognitive ability or relative cognitive decline may also themselves predict this decline^33^.

Testing PGSs as predictors of outcomes such as cognitive level and change is potentially useful and efficient. Researchers or clinicians can use a single source material—a participant’s DNA—to test their genetic propensity to a very wide range of risk and protective factors^34^. Therefore, instead of having to measure all the phenotypes that might confer risk to or protection of cognitive decline, it might be possible—to the extent that those phenotypes are heritable and have had a large, high-quality GWAS performed—to assess the genetic propensity to the phenotype and use that information to predict cognitive level and decline. The approach using PGSs, if successful, would also allow the retrospective testing of risk and protective factors in cohorts where DNA and longitudinal cognitive data are available but who were never tested for the risk or protective factors in question. Beyond these possible strengths, PGSs can even be used to assess propensity to a phenotype that is never expressed, such as liability to schizophrenia in a sample in which no one develops the illness^26^.

We selected fourteen PGSs based on, first, the relevant phenotype having been linked to cognitive decline in at least one previous study and, second, on there being a recent GWAS of that phenotype (Table S1 provides a list of references to the phenotypic studies, and to the respective GWASs). The PGSs in question were those for the following variables: educational attainment, the personality traits of neuroticism and conscientiousness, Alzheimer’s disease, Parkinson’s disease, schizophrenia, major depressive disorder, coronary artery disease, stroke, type 2 diabetes, smoking, height, body mass index, lung function, and grip strength. We tested the associations of each of these PGSs with the level (at age 70 years) and age-related slope (from age 70 to age 79 years) of general cognitive ability estimated from a battery of thirteen varied tests. We added a further analysis where we tested the association of the PGSs with change between a cognitive test taken at age 11 and age-70 general cognitive ability. We tested their predictive value individually, simultaneously, and— because the presence of the *APOE* e4 allele has previously been found to predict cognitive decline in this same cohort during almost the same period of life^5^—in models also including the *APOE* e4 status of the participants.

## Method

### Sample

The Lothian Birth Cohort 1936 (LBC1936) is an ongoing longitudinal study of older, community-dwelling individuals living mostly in the Edinburgh and Lothians area of Scotland, UK^35,36^. They were recruited on the basis of their having been part of the Scottish Mental Survey of 1947^37^, and have, to date, attended four testing waves: the first at age 69.54 years (SD = 0.83; *n* = 1,091; 543 females), the second at age 72.52 years (SD = 0.71; *n* = 866; 418 females), the third at age 76.25 years (SD = 0.68; *n* = 697; 337 females), and the fourth at age 79.32 (SD = 0.62; *n* = 550; 275 females). For simplicity, we will henceforth refer to the ages at each wave as 70, 73, 76, and 79 years, respectively. Ethical approval for the LBC1936 study came from the Multi-Centre Research Ethics Committee for Scotland (MREC/01/0/56; 07/MRE00/58) and the Lothian Research Ethics Committee (LREC/2003/2/29). All participants, who were volunteers and received no financial or other reward, completed a written consent form before any testing took place.

### Cognitive measures

In addition to completing the Moray House Test No. 12 at age 11 years^38^, which measures a variety of cognitive domains with an emphasis on verbal reasoning, the LBC1936 members completed a wide selection of cognitive tests at each of the later-life testing waves. Tests were administered identically at each occasion. Thirteen tests were used for the present analysis, covering the four broad cognitive domains described below.

#### Visuospatial ability

was measured using tests of pattern-based reasoning, recognition, and recall: the Matrix Reasoning and Block Design subtests of the Wechsler Adult Intelligence Scale, 3^rd^ UK Edition (WAIS-II^UK 39^), and the Spatial Span subtest of the Wechsler Memory Scale, 3^rd^ UK Edition (WMS-III^UK 40^; the score used here was an average of forwards and backwards spatial span).

#### Verbal memory

was measured using three tests of recall of new verbal information: the Logical Memory and Verbal Paired Associates subtests of the WMS-III^UK^ (both indicated by their total score), and the Digit Span backwards subtest of the WAIS-III^UK^.

#### Crystallized ability

was measured by three tests: the National Adult Reading Test (NART^41^), the Wechsler Test of Adult Reading (WTAR^42^) and a test of phonemic verbal fluency^43^. All three tests assessed prior verbal knowledge.

#### Processing speed

was measured using four tests tapping cognitive speed in a variety of ways. Two of the tests were pencil-and-paper “clerical” tasks: the Digit-Symbol Substitution and Symbol Search tasks from the WAIS-III^UK^. A third was a psychophysical measure of Inspection Time performed on a computer monitor (as described in ref.^44^). A fourth was a test of Choice Reaction Time, measured using the dedicated button-box described in ref.^45^. Note that, in each of the analyses, we reversed scores on the Choice Reaction Time test so that higher scores would indicate better cognitive performance.

### Genetic measures

The majority of participants provided blood samples at the age 70 wave that were used to extract DNA for the genetic analyses. To measure single-nucleotide polymorphisms (SNPs) we used the Illumina 610-Quadv1 whole-genome SNP array; measurements were completed at the Wellcome Trust Clinical Research Facility Genetics Core, Western General Hospital, Edinburgh. Polygenic scores (PGSs) were created using PRSice software^46^, with linkage-disequilibrium clumping parameters set to *r*^2^ > 0.25 over 250kb sliding windows. All PGSs were calculated using all SNPs from their respective GWAS (see Table S1 for all references); that is, we used an association threshold of *p* = 1.00. In four cases, we ran a new GWAS on data we had available from the UK Biobank sample (see Supplementary Method and Figures S1-S3). This was either because this resulted in a larger GWAS than the most recent published GWAS at the time, or because the LBC1936 participants were included in that most recent GWAS. In addition to the PGS analyses, each participant’s *APOE* e4 genotype was ascertained using TaqMan technology, also at the Wellcome Trust Clinical Research Facility Genetics Core. Since there were few carriers of two *APOE* e4 alleles (~2% of the sample), we categorised this variable as the binary presence (306 participants; ~30%) or absence (722 participants; ~70%) of any *APOE* e4 alleles.

### Statistical analysis and preregistration

In a set of preliminary analyses, we estimated whether each polygenic score was significantly associated with its “own” phenotype in the Lothian Birth Cohort. We selected phenotypes that were as closely-related as possible given the data we had available. The selected phenotypes were as follows. For the education PGS, we used years of education, reported at age 70. For the Neuroticism and Conscientiousness PGSs, Neuroticism and Conscientiousness were estimated using the NEO-FFI personality instrument^47^, completed at age 70. For the Alzheimer’s disease and Schizophrenia PGR, we used WAIS-III Block Design at age 70 (since this test provided an estimate of cognitive ability, which is impaired in both disorders, and no test of schizophrenia symptoms was available). For the major depressive disorder PGR, we used the score on the depression subscale of the Hospital Anxiety and Depression Scale, taken at age 70 (HADS^48^). For the coronary artery disease, stroke, and type 2 diabetes PGRs, we used self-reports of whether the participants had ever received a diagnosis of any of these conditions by age 70. For the smoking PGR, we used a self-report of whether the participant was a never-, ex-, or current smoker at age 70. For the height and BMI PGSs, we used the measurements of these traits taken by nurses at the age-70 testing wave. Finally, for the FEV1 and grip strength PGSs, we used the measurements of these physical functions taken at age 70 using a spirometer and a dynamometer, respectively.

The analyses described below were preregistered, except for the final one described below (including age-11 intelligence scores to estimate lifetime cognitive change), which, therefore, should be considered as exploratory. The time-stamped preregistration document, written after data from the fourth testing wave of LBC1936 were entered but before any of these data had been seen by any of the authors of this study, can be found at the following URL: https://osf.io/vyy4u/.

Before analysing any of the cognitive data, we used a parallel analysis, using the *psych* package for R^49^, to factor-analyse the PGSs, testing whether there was evidence for a general factor (as there is for cognitive tests). If evidence of such a general factor emerged, we planned to assess the value of this general factor as a predictor in the cognitive models.

We estimated a “factors of of curves” structural equation model to characterise cognitive levels and changes within older age. This involved estimating a latent growth curve model for each cognitive test, then factor-analysing the latent intercepts and latent slopes from these models. The model follows the same structure as that of ref.^5^, where we used data from the first three waves of the LBC1936 to examine predictors of cognitive change from age 70 to 76 years. The factor models for both levels and slopes were hierarchical, as shown in Figure 1: there were four domain-level factors estimated for both level and slope (Visuospatial ability, Verbal Memory, Crystallized ability, and Processing Speed), which were themselves factor-analysed to produce the general factors of cognitive level and slope.

**Figure 1.**
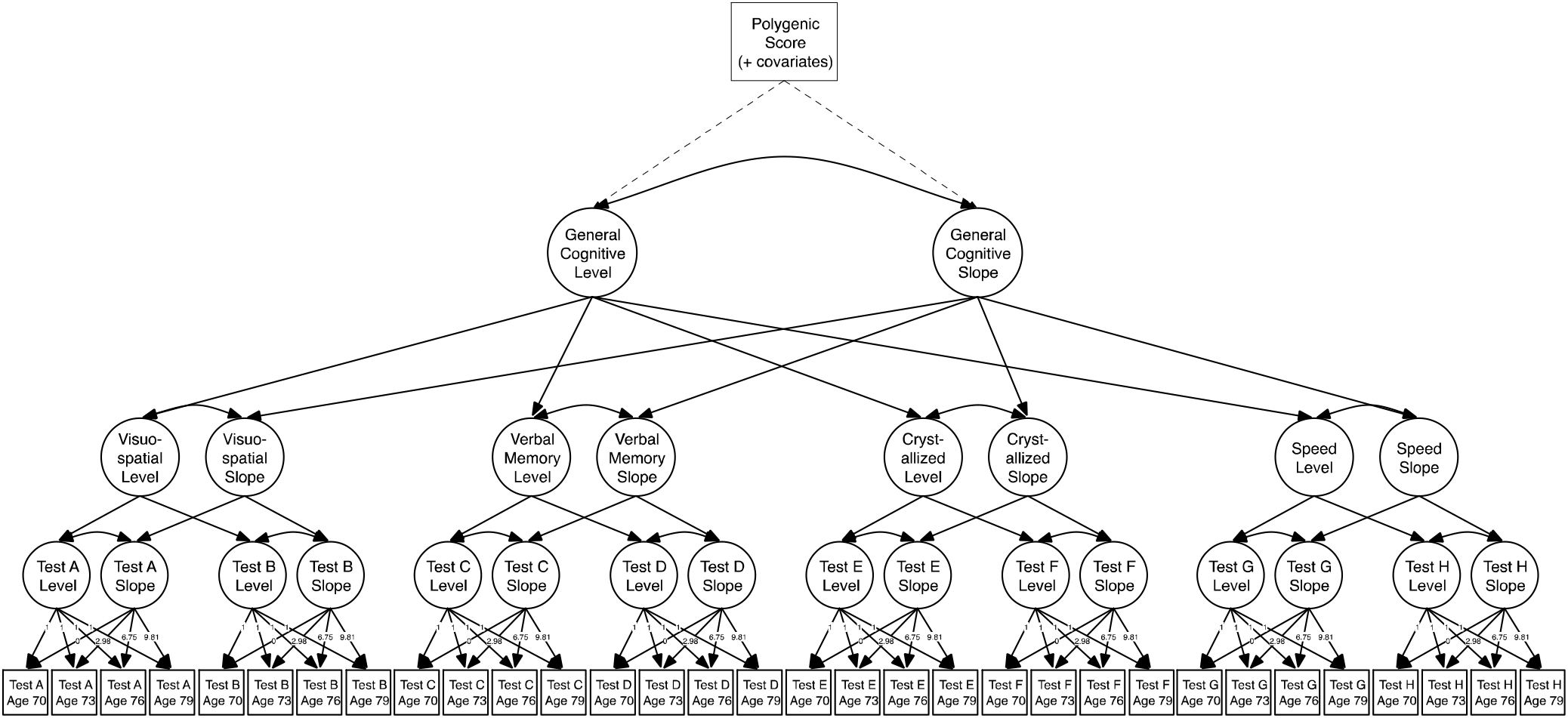
Simplified diagram of the structural equation model used to estimate the general cognitive level and the general cognitive slope. A latent growth curve was estimated across the four waves for each cognitive test (with the numbers showing the average length between each wave), and the levels and slopes were factor-analysed in a hierarchical model with four cognitive domains and a general factor. Note that, for illustrative purposes, not all tests are shown (the Speed domain had four tests and the other domains had three each; see Table 3). The main outcomes of interest—the association of each polygenic score with the general level and slope variables—are indicated by the dashed lines.

To estimate the mean change in each cognitive ability over time, we first ran a model with the raw cognitive scores. The models where we included the genetic predictors had the cognitive scores pre-residualised for age in days at the time of testing and for sex. The models with genetic predictors also included the four genetic principal component variables. Where possible, the analyses were run within the structural equation model (that is, the factors of curves model and the association with the predictors were estimated simultaneously). However, in some cases where the structural equation model would not converge, we extracted factor scores for the general factors and used linear regression to predict them from the genetic predictors. All such models are noted below.

We next ran the further analyses testing each PGS as a predictor alongside (that is, controlling for) *APOE* e4 status, to test for incremental predictive validity over this well-established risk factor. We did this only for PGSs that had shown a significant relation to the relevant cognitive outcome in the main analysis. Then, we included all the PGSs together as predictors in a single model, assessing their incremental validity over one another. For a final analysis, which was not part of the preregistration, we tested whether each genetic predictor (the PGSs and *APOE* e4) was associated with age-70 general cognitive ability after correcting for the age-11 intelligence measure. In this way, we were able to test whether each measure was predictive of cognitive change across most of the life course (that is, between ages 11 and 70).

The LBC1936 participants had a homogeneous, White European background, and thus we did not expect population stratification to have a strong influence on the results. Nevertheless, for all analyses involving relating the PGSs (and *APOE* e4) to phenotypes, we included four SNP principal components (multidimensional scaling components) as covariates. We used the False Discovery Rate correction^50^ to adjust *p*-values for multiple comparisons across the 15 predictors (14 PGSs plus *APOE* e4). Structural equation modelling was performed using Mplus v7.3^51^, and used full-information maximum likelihood estimation to use all of the available data in each model. All other analyses were run in R^52^.

## Results

### Preliminary polygenic score analyses

Thirteen of the fifteen PGSs were significantly associated with their “own” related phenotype (see Table 1) in the LBC1936 sample. These significant relations ranged from explaining 1.00% of the variance (for both the grip strength and the major depressive disorder PGSs in grip strength and HADS depression score respectively), to the PGS for BMI explaining 10.69% of the variance in BMI. Two of the scores were not significantly related to the outcome phenotype: the PGS for Alzheimer’s disease was not significantly related to cognitive ability (as measured by the Block Design test; explaining 0.02% of the variance, *p* = .64), and the PGS for stroke was not significantly related to self-reported stroke (explaining 0.64% of the variance, *p* = .15).

**Table 1.** Variance explained by each polygenic profile score in relevant Lothian Birth Cohort 1936 outcome variables.

| Polygenic Profile Score | Outcome Variable | Std. $\beta$ | SE | *p*-value | % Variance Explained |
| --- | --- | --- | --- | --- | --- |
| Educational attainment | Years of education | **.22** | **.03** | **$6.09 \times 10^{-12}$** | **4.64%** |
| Neuroticism | NEO-PI-R Neuroticism | **.18** | **.04** | **$1.85 \times 10^{-06}$** | **2.58%** |
| Conscientiousness | NEO-PI-R Conscientiousness | **.08** | **.03** | **.01** | **0.69%** |
| Alzheimer's disease | Block Design | -.02 | .03 | .63 | 0.02% |
| Schizophrenia | Block Design | **-.12** | **.03** | **$4.57 \times 10^{-04}$** | **1.23%** |
| Major depressive disorder | HADS depression score | **.10** | **.03** | **.002** | **1.00%** |
| Coronary artery disease | Cardiovascular disease<sup>†</sup> (24.5%) | **.27** | **.08** | **$3.94 \times 10^{-04}$** | **1.89%** |
| Stroke | Stroke<sup>†</sup> (5.0%) | .21 | .15 | .15 | 0.64% |
| Type 2 diabetes | Type 2 diabetes<sup>†</sup> (8.6%) | **.53** | **.13** | **$2.40 \times 10^{-05}$** | **4.17%** |
| Smoking | Smoking | **.15** | **.03** | **$5.41 \times 10^{-06}$** | **2.05%** |
| Height | Height | **.28** | **.03** | **$2.27 \times 10^{-16}$** | **6.53%** |
| BMI | BMI | **.33** | **.03** | **~.00** | **10.69%** |
| FEV<sub>1</sub> | FEV<sub>1</sub> | **.19** | **.03** | **$3.00 \times 10^{-13}$** | **5.21%** |
| Grip strength | Grip strength | **.06** | **.02** | **.002** | **1.00%** |

*Note.* Standardized $\beta$ s, SEs, and *p*-values are from general linear (continuous outcomes) or generalized linear (categorical outcomes, indicated with the <sup>†</sup> symbol and with the percentage of the sample who reported having, or having had, that condition at or by age 70) regression models adjusting for age at the time of measuring/reporting the outcome variable, sex, and four multidimensional scaling components. All significant *p*-values remained significant after False Discovery Rate correction. For continuous outcome variables, the % variance explained is derived from the partial $R^2$ . For categorical outcome variables, the % variance explained is derived from the Nagelkerke's $R^2$ . Rows in bold survived false discovery rate correction for multiple testing. References for the GWAS source of each polygenic score can be found in Table S1.

A correlation matrix of the relations among each of the PGSs is displayed in Table 2. The PGS for education showed the highest number of significant relations to the other PGSs, being related to nine of the other fourteen scores; this mostly consisted of education-linked variants being positively associated with variants linked to better health (broadly consistent with evidence from genetic correlations; see e.g. ref^53^). The correlation sizes were generally low: the strongest relation between any of the PGSs was that between the education and smoking PGSs, estimated at *r* = .25. As planned in the preregistration, we ran a parallel analysis of the fifteen PGSs using the *psych* package for R: this indicated that there were six factors in the data, with no evidence for a strong “general” factor. Therefore, we did not use any of these factors in the analyses below.

**Table 2.** Pearson correlations between polygenic scores in the Lothian Birth Cohort 1936.

| Polygenic Score | 1. | 2. | 3. | 4. | 5. | 6. | 7. | 8. | 9. | 10. | 11. | 12. | 13. |
| --- | --- | --- | --- | --- | --- | --- | --- | --- | --- | --- | --- | --- | --- |
| 1. Education | - |  |  |  |  |  |  |  |  |  |  |  |  |
| 2. Neuroticism | -.11*** | - |  |  |  |  |  |  |  |  |  |  |  |
| 3. Conscientiousness | -.02 | -.04 | - |  |  |  |  |  |  |  |  |  |  |
| 4. Alzheimer's disease | -.11*** | .01 | .0004 | - |  |  |  |  |  |  |  |  |  |
| 5. Schizophrenia | .004 | .08** | -.03* | -.02 | - |  |  |  |  |  |  |  |  |
| 6. Major depressive disorder | -.06* | .18*** | -.04 | .04 | .17*** | - |  |  |  |  |  |  |  |
| 7. Coronary artery disease | -.13*** | .004 | -.03 | -.003 | .06* | .02 | - |  |  |  |  |  |  |
| 8. Stroke | -.10** | .01 | -.02 | -.01 | .05 | .09** | .11*** | - |  |  |  |  |  |
| 9. Type 2 diabetes | -.03 | .02 | .02 | .02 | .08* | -.06 | .12*** | -.05 | - |  |  |  |  |
| 10. Smoking | -.25*** | .06* | -.01 | .03 | .12*** | .04 | .08** | .03 | .02 | - |  |  |  |
| 11. Height | .09** | -.04 | .02 | -.10*** | -.15*** | -.10** | -.03 | .07* | -.23*** | -.04 | - |  |  |
| 12. BMI | -.16*** | -.05 | -.04 | .03 | .04 | -.003 | .07* | .04 | .16*** | .14*** | -.11*** | - |  |
| 13. FEV<sub>1</sub> | .13*** | .09* | -.07* | .03 | .13*** | .06* | -.01 | -.05 | -.04 | -.13*** | -.07* | -.08* | - |
| 14. Grip strength | .03 | -.12*** | .05 | -.08 | .03 | -.09** | .03 | -.01 | -.005 | -.03 | -.04 | .04 | .15*** |

Note: $n = 1,005$ for all correlations. \* $p < .05$ , \*\* $p < .01$ , \*\*\* $p < .001$ .

### Cognitive levels and slopes

Descriptive statistics for each of the cognitive tests at each wave, and a full correlation matrix for the cognitive tests, are provided in Tables S2 and S3 in the Supplementary Materials, respectively. The longitudinal changes, estimated from the structural equation models, for each cognitive test score are shown in Table 3. All but two of the tests showed statistically significant declines over time, with the largest per-year declines being seen in the processing speed tests. The two tests showing no significant age-related change were the NART and verbal fluency, both of which were in the category of “crystallized” tests and were thus expected to show less decline with age. The trajectory of each test with age is illustrated in Figure 2, for the model-implied trajectory as well as the change in the raw data for comparison (see Figure S4 for an alternative way of visualizing the data, showing each data point). In all but one case the model-implied trajectory was the same in sign as that in the raw data, though some slopes were different in magnitude. For verbal paired associates, the magnitude was reversed (it became negative in the model). These changes are to be expected given the use of FIML estimation in the structural equation model: several previous methodological investigations have shown that the choice of missing-data technique can influence results^54,55^. Foster et al.^56^ recommend that the technique is chosen on the basis of its assumptions being plausibly fit to a dataset; on the basis of a previous analysis where few of the LBC1936 sample’s predictor variables could predict dropout from age 70 to age 76^5^, we conclude that the assumption of “missing at random”, the basis for the use of FIML, was justified here.

**Figure 2.**
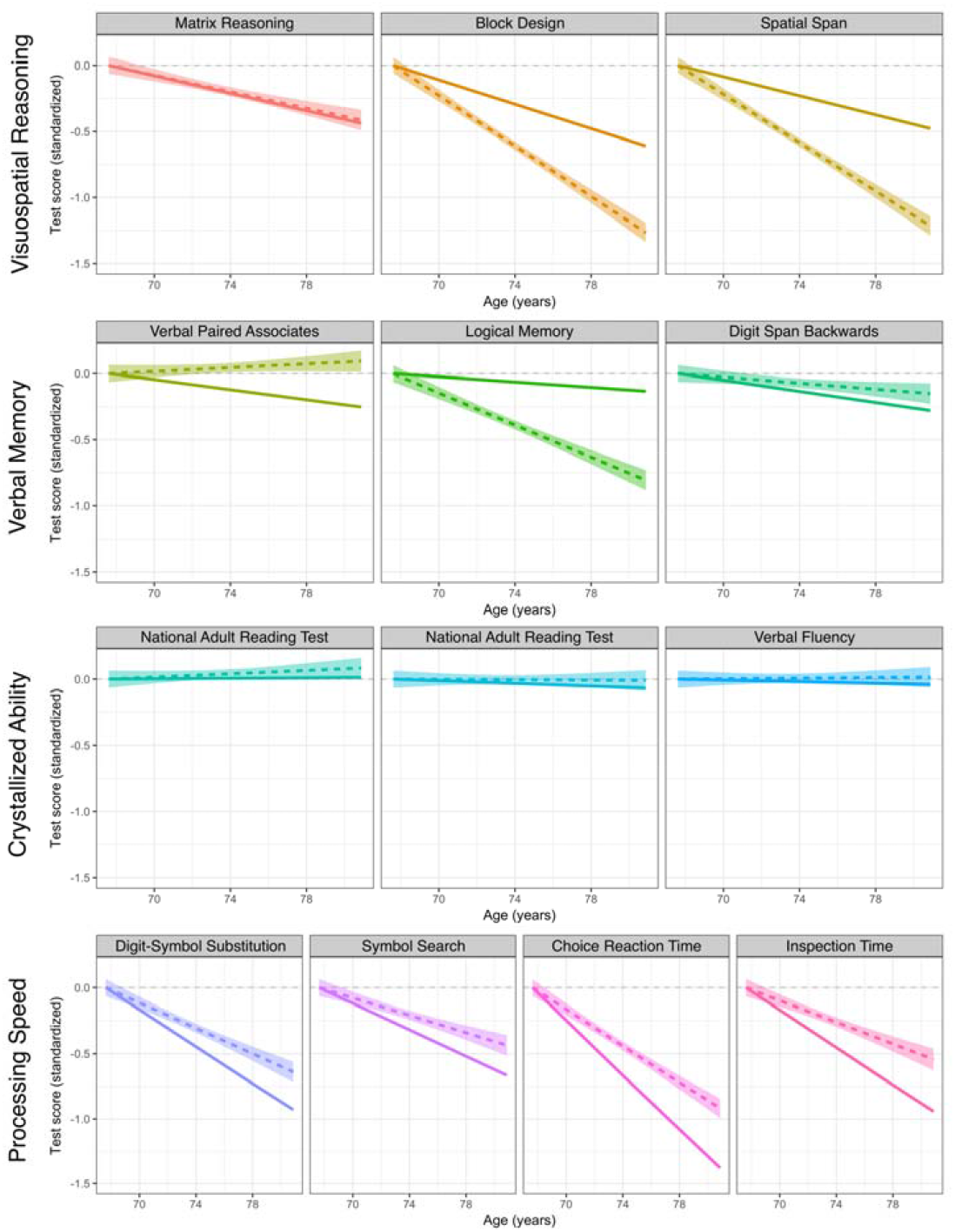
Standardized linear trajectories of each cognitive test with age. Intercepts (at the youngest age) are set to zero for comparative purposes. The horizontal dotted line indicates zero. The coloured solid line is the model-implied trajectory (using full-information maximum likelihood estimation); the coloured dotted line is the regression line through the raw data (with shaded 95% confidence interval).

**Table 3.** Estimates of the linear slope of each cognitive test. Illustrations of each trajectory are shown in Figure 2.

| Domain | Cognitive test<br>(max. score) | Mean<br>(SD) at<br>age-70<br>baseline | Mean<br>raw<br>change<br>per<br>year | SE of<br>raw<br>change | *p*-value | Mean<br>no. of<br>SDs<br>change<br>per year |
| --- | --- | --- | --- | --- | --- | --- |
| Visuospatial<br>ability | Matrix<br>reasoning | 13.49<br>(5.13) | −0.133 | 0.016 | $6.12 \times 10^{-16}$ | −0.033 |
| | Block design | 33.79<br>(10.32) | −0.423 | 0.029 | $1.30 \times 10^{-49}$ | −0.046 |
| | Spatial span | 7.36<br>(1.42) | −0.038 | 0.005 | $1.27 \times 10^{-15}$ | −0.036 |
| Verbal<br>memory | Logical<br>memory | 71.46<br>(17.96) | −0.150 | 0.072 | .038 | −0.010 |
| | Verbal paired<br>associates | 26.44<br>(9.13) | −0.156 | 0.033 | $3.00 \times 10^{-06}$ | −0.019 |
| | Digit span<br>backwards | 7.73<br>(2.26) | −0.038 | 0.007 | $1.72 \times 10^{-07}$ | −0.021 |
| Crystallized<br>ability | NART | 34.48<br>(8.15) | 0.012 | 0.013 | .379 | 0.001 |
|  | WTAR | 41.02<br>(7.17) | −0.034 | 0.012 | .003 | −0.005 |
|  | Verbal fluency | 42.42<br>(12.54) | −0.032 | 0.035 | .358 | −0.003 |
| Processing<br>speed | Digit-symbol<br>substitution | 56.60<br>(12.93) | −0.833 | 0.038 | $5.44 \times 10^{-105}$ | −0.070 |
| | Symbol search | 24.71<br>(6.39) | −0.258 | 0.023 | $1.50 \times 10^{-28}$ | −0.050 |
| | Inspection time | 112.14<br>(11.00) | −0.595 | 0.049 | $2.26 \times 10^{-34}$ | −0.071 |
| | Choice reaction<br>time (ms) | 64.21<br>(0.09) | −0.008 | $3.57 \times 10^{-04}$ | $1.86 \times 10^{-108}$ | −0.104 |

*Note:* Estimates of SD change are model-implied, using full-information maximum likelihood estimation, and thus may not precisely correspond to the raw SDs.

We then factor-analysed the levels and slopes of all the tests, as described above and shown in Figure 1. The baseline model (with the tests corrected for age within the wave and sex, but with no polygenic score predictors) showed excellent fit to the data, according to multiple fit indices: χ^2^(1298) = 2446.16, *p* < .001; Root Mean Square Error of Approximation = 0.028; Comparative Fit Index = 0.968; Tucker-Lewis Index = 0.967; Standardized Root Mean Square Residual = 0.057. The full parameters for this model are shown in Figure S5 in the Supplementary Materials. Squaring the product of each test’s loading on its domain and the domain’s loading on the general factor provided a proportion of variance in each test explained by the general factor; averaging these figures showed that the general factor of cognitive level explained 40.9% of the variance in performance across all the tests, and the general factor of cognitive slope explained 69.7% of the variance in cognitive change between ages 70 and 79.

### Polygenic score prediction of general cognitive level and slope

The associations of each PGS and *APOE* e4 with the age-70 level and age 70-to-79 slope of general cognitive ability are shown in Table 4 and Figure 3. Of the fifteen PGSs, six were statistically significantly associated with cognitive level at age 70: higher PGSs for education and height were associated with higher general cognitive levels, and PGSs for schizophrenia, coronary artery disease, smoking, and BMI were associated with lower general cognitive levels. All effect sizes for these significant effects were modest: the standardized betas ranged from –.178 (for smoking) to .264 (for education), the p-values that remained significant after multiple-comparisons correction ranged from 1.54×10^-15^ to .002. *APOE* e4 status was only nominally significantly associated with level: its association became nonsignificant after correction for multiple comparisons.

**Figure 3.**
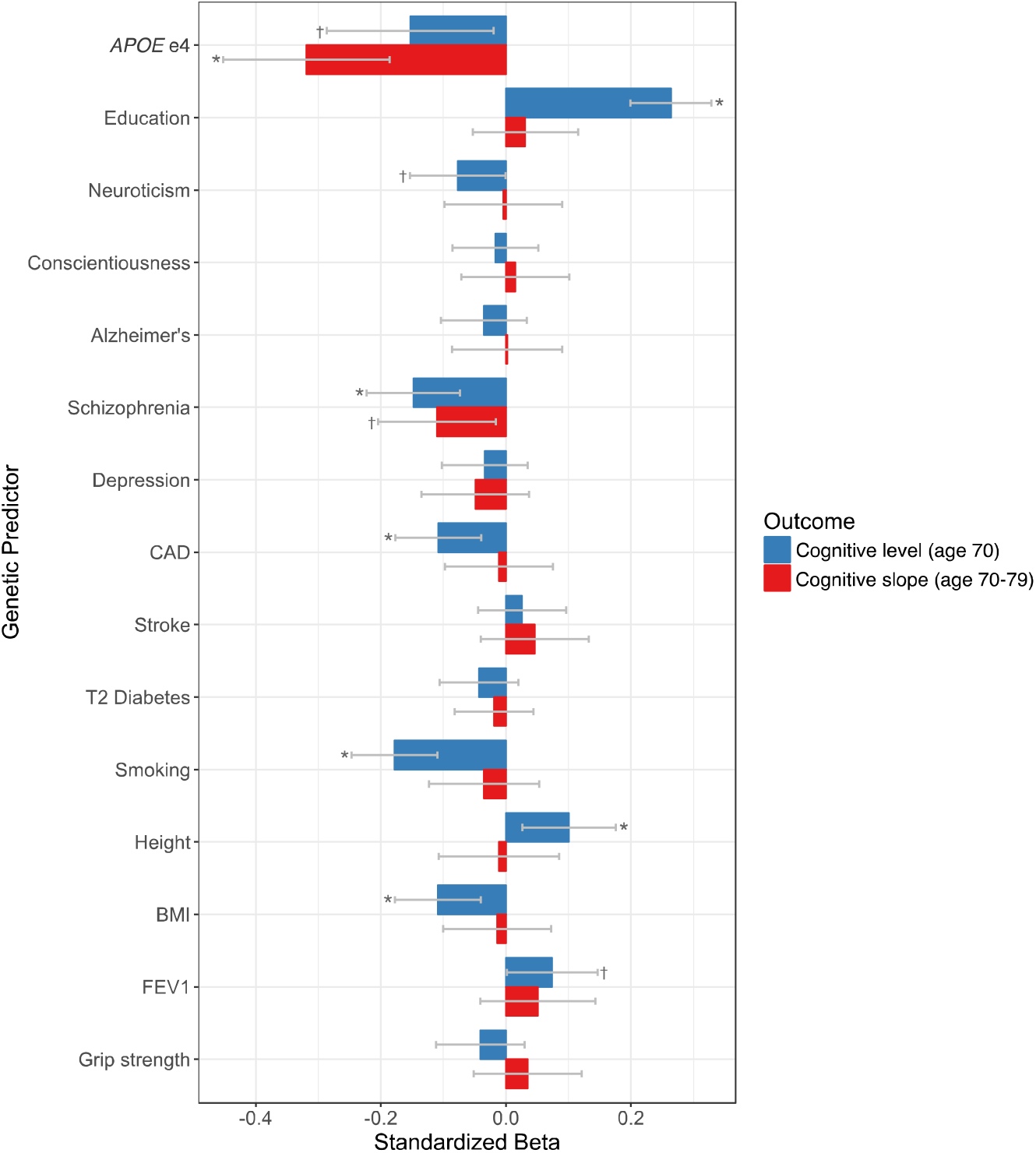
Associations of *APOE* e4 status and each polygenic score with cognitive level (age 70) and cognitive decline (age 70 to 79). * = statistically significant after false-discovery rate correction. † = nominally significant, but no longer significant after false-discovery rate correction. Note that the effect for *APOE* e4 is standardized only with respect to the outcome (with a dichotomous predictor); all other effect sizes are standardized with respect to both the outcome and the predictor.

**Table 4.** Associations of each polygenic profile score, and *APOE* e4 status, with general cognitive level (age 70) and slope (age 70-79) in individual-predictor models.

| Genetic variable | Association with baseline *g* |  |  | Association with *g* slope |  |  |
| --- | --- | --- | --- | --- | --- | --- |
| | Std. $\beta$ | SE | *p*-value | Std. $\beta$ | SE | *p*-value |
| *APOE* e4 | -.153 | .068 | .025 | **-.319** | **.068** | **$3.44 \times 10^{-06}$** |
| Education | **.264** | **.033** | **$1.54 \times 10^{-15}$** | .031 | .043 | .474 |
| Neuroticism | -.077 | .039 | .047 | -.004 | .048 | .929 |
| Conscientiousness | -.017 | .035 | .634 | .015 | .044 | .726 |
| Alzheimer's disease | -.035 | .035 | .316 | .002 | .045 | .957 |
| Schizophrenia | **-.148** | **.038** | **$1.07 \times 10^{-04}$** | -.110 | .048 | .022 |
| Major depressive disorder | -.034 | .035 | .342 | -.049 | .044 | .267 |
| Coronary artery disease | **-.108** | **.035** | **.002** | -.011 | .044 | .808 |
| Stroke | .026 | .036 | .463 | .046 | .044 | .295 |
| Type 2 diabetes | -.043 | .032 | .171 | -.019 | .032 | .550 |
| Smoking | **-.178** | **.035** | **$3.66 \times 10^{-07}$** | -.035 | .045 | .438 |
| Height | **.101** | **.038** | **.008** | -.011 | .049 | .826 |
| BMI | **-.109** | **.035** | **.002** | -.014 | .044 | .746 |
| FEV<sub>1</sub> | .074 | .037 | .048 | .051 | .047 | .277 |
| Grip strength | -.041 | .036 | .255 | .035 | .044 | .425 |

Note: All estimates come from hierarchical latent growth curve structural equation models. Associations are corrected for age at cognitive testing, sex, and four multidimensional scaling components. Bold values are those that were statistically significant after false discovery rate correction for multiple testing. Note that the results for *APOE* e4 and Type 2 diabetes were estimated using extracted factor scores in linear regression models; all other effect sizes were estimated within the structural equation models themselves.

Only one of the PGSs, that for schizophrenia, was a significant (negative) predictor of general cognitive slope between ages 70 and 79, but this did not survive multiple-comparisons correction. *APOE* e4 status did, however, was a significant predictor of general cognitive decline—those with one or two *APOE* e4 alleles had significantly steeper general cognitive decline than those who had none—and this association survived multiple-testing correction.

We next tested whether the scores that had shown significant relations to the cognitive outcomes in the initial analysis were still statistically significantly related after including *APOE* e4 as a separate predictor. Results are shown in Table S4. In all cases, the polygenic score predictor remained significant after adjusting for *APOE* e4 status, with slightly attenuated effect sizes: that is, their associations seemed to be largely independent of any relation with *APOE* e4.

Next we ran the simultaneous-predictor models, including all PGSs predicting general cognitive level, and (in a separate model) general cognitive slope. For general cognitive level at age 70, only the significant associations with the PGSs for education (*β* = .195) and schizophrenia (*β* = –. 126) remained (see Table S5 for full results). As in the initial analysis, none of the PGSs were significantly related to general cognitive slope over time after correction for multiple comparisons.

### Additional (non-preregistered) analyses

We ran the analyses predicting general cognitive level at age 70 adjusted for age-11 cognitive ability – that is, predicting cognitive change across most of the life course. The basic cognitive model is shown in Figure S6. Note that age 11 cognitive ability was related very strongly to age 70 general cognitive level (*β* = .814, SE =.015, *p* = ~.00; see ref.^5^). Results from the genetic analyses are shown in Table 5, along with correlations between each of the genetic predictors and age-11 cognitive ability itself. Five of the sixteen PGSs were significantly associated with intelligence score at age 11 after multiple-comparisons correction: education, schizophrenia, coronary artery disease, smoking, and BMI. The association between lifetime cognitive change and the PGS for education was significant (and survived multiple-testing correction), with a standardized effect size of *β* = .102: those with a higher PGS for education saw relatively less cognitive change across the lifespan. The other PGSs were not related significantly to the lifetime cognitive change variable.

**Table 5.** Associations of each polygenic score and *APOE* e4 status with age 11 intelligence and lifetime cognitive change (age 70 general intelligence (*g*) adjusted for age 11 intelligence)

| Genetic variable | Association with age 11 intelligence |  |  | Association with age 70 *g*, adjusted for age 11 intelligence |  |  |
| --- | --- | --- | --- | --- | --- | --- |
| | Std. $\beta$ | SE | *p*-value | Std. $\beta$ | SE | *p*-value |
| *APOE* e4 | .024 | .074 | .740 | −.080 | .055 | .141 |
| Education | **.236** | **.032** | **6.13×10<sup>−13</sup>** | **.102** | **.025** | **5.19×10<sup>−05</sup>** |
| Neuroticism | −.032 | .037 | .377 | −.016 | .027 | .563 |
| Conscientiousness | .010 | .033 | .741 | −.171 | .119 | .153 |
| Alzheimer's disease | −.011 | .033 | .731 | −.026 | .025 | .282 |
| Schizophrenia | **−.136** | **.036** | **1.85×10<sup>−04</sup>** | −.021 | .028 | .446 |
| Major depressive disorder | −.028 | .033 | .400 | .028 | .025 | .256 |
| Coronary artery disease | **−.098** | **.032** | **.003** | −.039 | .025 | .116 |
| Stroke | −.003 | .033 | .927 | .052 | .025 | .038 |
| Type 2 diabetes | −.024 | .035 | .489 | −.006 | .025 | .818 |
| Smoking | **−.187** | **.033** | **2.60×10<sup>−08</sup>** | −.023 | .026 | .369 |
| Height | .053 | .036 | .142 | .030 | .027 | .270 |
| BMI | **−.128** | .033 | **1.44×10<sup>−04</sup>** | −.012 | .025 | .636 |
| FEV<sub>1</sub> | .055 | .035 | .113 | .023 | .026 | .380 |
| Grip strength | −.030 | .033 | .362 | −.002 | .025 | .927 |

*Note:* Lifetime change come from a hierarchical latent model of general cognitive ability, comprising tests taken at age 70 (Figure S3), with the *g-factor* adjusted for age 11 intelligence. Rows in bold are effects that survived (per-column) false discovery rate correction for multiple testing. All models included four principal components to adjust for population stratification.

We did not expect that the Alzheimer’s disease PGS would make such a poor (near-zero; see Table 4) prediction of general cognitive slope, especially when *APOE* e4 status had made a significant prediction, and SNPs associated with *APOE* e4 featured prominently in the Alzheimer’s GWAS results from which the PGR was calculated^24^. We tested whether this was due to our use of the *p* = 1.00 threshold when calculating the PGS—it is possible that inclusion of the large number of less- and non-significant SNPs overwhelmed the signal from *APOE* e4-linked loci—by recalculating the PGS at a more stringent threshold—*p* = .01—and re-running the relevant analyses. As would be expected, the more stringent PGS was had a somewhat higher (point-biserial) correlation with *APOE* e4 status (r(964) = .251, *p* = 2.43 ×10^-15^) than did the PGS with the *p* = 1.00 threshold (r(963) = .097, *p* = .002). The more stringent Alzheimer’s PGS also made a significant prediction of the slope of general cognitive decline (*β* = –.128, SE = .043, *p* = .003), though not its initial level (*β* = -.016, SE = 0.035, *p* = .648).

For comparison, we also re-calculated the other PGSs at the *p* = .01 threshold. The results for all *p* = .01 PGSs are shown in Table S6. Most results were similar: unlike the Alzheimer’s result reported above, all other non-significant associations with general cognitive slope remained non-significant with this new threshold. The predictions for cognitive level generally had similar effect sizes, though the association between level and coronary artery disease was no longer significant, while the nominally-significant associations for the height and BMI PGSs did not survive multiple comparisons correction.

## Discussion

The study reported here was an attempt to predict variation in cognitive ageing using polygenic scores. All the PGSs were chosen *a priori* to index genetic variants associated with key ageing-relevant traits and conditions. Although some of these PGSs were statistically significantly associated with the baseline cognitive level at age 70, and the education PGS was associated with cognitive change from age 11 to age 70, none of the PGSs was significantly associated with the gradient of the general cognitive slope between age 70 and 79 – a general cognitive decline variable that explained over two-thirds of the slope variance across all thirteen tests). The presence of the *APOE* e4 allele, on the other hand, could make significant predictions of cognitive decline: having either one or two alleles, compared to zero, explained around 10% of the variance in the slope of cognitive decline. None of the PGSs approached this level of effect size. Below, we discuss some of the implications, strengths, and limitations of the study.

Overall, the results were similar to results from studies attempting to use phenotypic data to predict cognitive decline (e.g. attempts in this same sample over a shorter period of change^5^): cognitive levels could be predicted far more easily than cognitive slopes. It should be noted that some of the PGS correlations with cognitive level were similar in effect size to those using the actual phenotypes themselves, examined in previous studies. For example, the authors in ref.^5^ found a correlation between measured BMI and age-70 cognitive level of – .111 (see their Table S5); here, the PGS for BMI had a correlation with the same latent variable of –.109. It may be that any genetic effects indexed by the PGS occur at different points in the life course. For instance, variants linked to education and BMI—two PGSs that were significantly associated with baseline cognitive level, but not slope—may have their effects on cognitive ability during childhood or early adulthood, whereas effects of *APOE* e4—which significantly predicted slope, but not level—may appear only within older age. In one GWAS of general cognitive ability^57^, the effect of *APOE* e4 was particularly pronounced at older ages, consistent with these findings (see ref.^58^ for a longitudinal analysis in an even older sample). Note that any PGS associations reported herein were independent of *APOE* e4, as per our simultaneous regression analysis including them both as predictors.

The result that the residual of cognitive ability between age 11 and age 70—an index of lifetime cognitive change—was statistically significantly predicted by the education PGS also implies that education-linked variants are related to changes in cognitive abilities before old age (specifically, before age 70). These findings are of relevance to the concept of “cognitive reserve”^13^: they imply that researchers who find links between early-life education and later-life cognitive abilities (or cognitive change) should take into account the fact that some of the effect may come from a genetic propensity to better educational attainment, and not the educational attainment itself.

Given the general difficulty of finding significant predictors of cognitive decline, it may simply take a higher-powered study, with more participants, longer follow-up times, and additional waves, to detect genetic effects. On the other hand, as was noted above, larger GWAS studies have tended to produce PGSs with better predictive validity^27^, and this is likely also the case here. Another reason for the lack of prediction may be because the phenotypes linked to the PGSs do not themselves reliably predict cognitive decline. The most recent systematic review^10^ noted that much of the evidence in this sphere is weak; we do not have a strong, multi-study evidence base for many of the phenotypic—let alone genetic— predictors of cognitive decline discussed here. It is also possible that genetic propensities interact with environments in ways that improve or worsen the cognitive ageing trajectory. Note that there is no existing large, well-powered GWAS of cognitive decline in particular, since there are too few samples with the relevant variables for such a GWAS to be run. Should such a well-powered study appear in future, we would expect to derive a PGS that would predict cognitive decline in separate samples.

The case of the Alzheimer’s PGS warrants further consideration. As noted above, we chose to calculate all PGSs at the most liberal, whole-genome threshold *(p* = 1.00), including the effects of all SNPs, rather than a more restricted set. The choice of just one single threshold was to avoid the overfitting that often comes with choosing many thresholds and reporting the one with the highest association with the outcome trait of interest. However, the fact that the prediction of cognitive decline by *APOE* e4 status was so much larger than that of the Alzheimer’s PGS, which itself contains the effect of (SNPs in linkage disequilibrium with) *APOE* e4, suggested that the effects of the other SNPs—less significantly or not significantly related to cognitive decline—included in the PGS were overpowering the *APOE* e4-linked signal. This appeared to the be the case: the use of a more conservative threshold improved the predictive validity. This was not the case for the education PGS, however: it may be that for traits that have even higher levels of polygenicity, for which there are no large-effect variants such as *APOE* e4 for Alzheimer’s, the PGS threshold matters less. As we noted above, larger GWASs of Alzheimer’s disease will produce summary data with more signals from non-APOE-e4-linked variants, and these should be tested for their association with normal-range cognitive ability and cognitive decline at different thresholds (note that a previous study^59^ found no relation between a PGS calculated from an older Alzheimer’s GWAS and cognitive abilities in this same sample; see also ref.^60^ for an example of a combination of *APOE* and PGS variables). Alternatively, performing a permutation test as described in ref.^46^ (Supplementary Note 4) would allow the calculation of an empirical p-value threshold, potentially allowing for a better prediction than we had with our across-the-board use of the *p* = 1.00 threshold. Generally, however, there are as yet no hard-and-fast rules for the use of PGSs by researchers who wish to maximise their predictive ability but are concerned about multiple-comparisons testing; ref.^60^ provides some recommendations.

Another strategy to improve prediction would be to follow the approach of ref.^61^, where the authors took a much larger set of eighty-one PGSs and used them as predictors of the levels of various traits including cognitive ability and BMI (they used a sample of younger individuals and thus could not examine cognitive decline). They ran their analysis using penalized regression (specifically the Elastic Net^62^), which allowed the large number of PGS variables to be reduced to the best set of predictors, and allowed them to explain a somewhat larger proportion of the variance in the traits. Such a “hypothesis-free” approach, using algorithmic selection from many predictors rather than manually choosing those predictors on the basis of pre-existing links to cognitive decline, as we did here, may be a more fruitful approach.

## Strengths and limitations

The LBC1936 sample is a rare dataset in that it is a narrow-age sample that covers nearly an entire decade within older age, with thirteen high-quality cognitive tests measured repeatedly and identically; moreover, and highly unusually for an ageing study, it has well-validated cognitive test data from age 11. Using longitudinal structural equation modelling, we estimated general factors of cognitive level and slope that removed any measurement error associated with the individual tests and produced an index of overall cognitive ability and its ageing. Overall, this was a powerful way to assess cognitive decline, and we used a variety of PGSs from varied traits and disorders in an attempt to predict that decline’s variance. However, there are a number of limitations to the study.

The self-selecting nature of the LBC1936 participants may have biased our results. That is, the participants were generally healthy, independently-living older adults and—by virtue of the fact they were able and interested to attend the initial testing—were likely substantially healthier and more intellectually engaged than the average person of their age. Non-random dropout, a problem for most longitudinal studies of ageing, compounds this issue: individuals who remained in the study across the four waves were healthier on average than those who dropped out. We thus probably missed individuals with the poorest health and, consequently, with the greatest degree of cognitive decline. This limitation—a restriction of range to the higher end of the health distribution—may have contributed to our lack of ability to predict cognitive decline in our sample.

There may also be additional complexities in the cognitive decline paths that we did not consider here, since we focused on the relatively simple linear average trajectories. For example, there may be nonlinear trajectories, or multiple latent trajectories (see e.g. ref.^63^) within the dataset that, if analysed in future, may reveal differential relations to the genetic predictors. Finally, in a few cases, the complexity of the structural equation models meant that convergence could not be achieved without fixing some of the paths that were intended to be freely-estimated (see Figure S2), and in some cases without extracting factor scores and using them in linear regression models instead of estimating all relations within latent-variable space (i.e. simultaneously to the estimation of the relevant structural equation model). We would not estimate that these issues would have changed our results to a great extent, but the latter practice (extracting factor scores) may have led to slight alterations to some of the standard errors we reported.

## Conclusions

A key goal of cognitive ageing research is to be able to predict who will experience steeper general cognitive decline. In this study of a high-quality, longitudinal dataset, we attempted to do so using a panel of polygenic scores, but were broadly unsuccessful: despite statistically significant associations of several polygenic scores (those for education, schizophrenia, coronary artery disease, smoking, height, and BMI) with general cognitive level at baseline, and the relation of the education PGS to cognitive change between age 11 and age 70, none of the scores predicted subsequent cognitive decline across the eighth decade of life in the pre-registered analysis. Future, larger GWAS studies might furnish us with summary statistics to produce more predictive PGSs, and different analytic methods might increase the predictive value of those we already have. Given that it is possible to formulate so many PGSs from only DNA, the approach retains its potential as an efficient and protean source of predictors. For the time being, however, the present study’s findings suggest that researchers interested in genetic prediction of longitudinal variability in cognitive ageing will derive more value from established predictors such as *APOE* e4 than newer methods such as polygenic scores.

## Acknowledgements

We thank the Lothian Birth Cohort 1936 members for their participation in the study, and the research team for the collection of the data. We are grateful to the Scottish Council for Research in Education for access to the data from the Scottish Mental Survey of 1947. The Lothian Birth Cohort 1936 data collection was supported by Age UK (Disconnected Mind programme grant). The work was undertaken in The University of Edinburgh Centre for Cognitive Ageing and Cognitive Epidemiology, part of the cross-UK Research Council Lifelong Health and Wellbeing Initiative (MR/K026992/1). Additional support from the Medical Research Council and the Biotechnology and Biological Sciences Research Council is greatly appreciated.

None of the authors report any relevant conflicts of interest.

